# Dynamic Ontogeny of Auditory Lateralization in the Zebra Finch

**DOI:** 10.1101/2025.03.30.646178

**Authors:** Basilio Furest Cataldo, Paige Dadika, David S. Vicario

## Abstract

Functional lateralization is a ubiquitous trait in the animal kingdom and represents a general and conserved mechanism of the central nervous system. Lateralized processes observed in adult sensory cortices emerge as a function of development and experience, e.g. speech-processing in humans and processing of conspecific vocalizations in songbirds. Adult Zebra finches (ZFs), a species of songbird, exhibit right-lateralized activity in the higher auditory region *caudomedial nidopallium* (NCM) which depends on normal rearing conditions; auditory deprivation leads to atypical bilateral responses to conspecific song. Here, we investigate the ontogenetic timeline of auditory lateralization and the lasting effects of auditory rearing-environment in the developing ZF (40-120 days post-hatch, phd). ZFs were raised in one of three acoustic environments: 1) a tutor-playback driven paradigm, 2) chronic exposure to a ZF aviary recording (no tutor), and 3) chronic exposure to a canary aviary recording (no tutor). By longitudinally tracking lateralized auditory evoked potentials at the level of the dura, we show that adult right-lateralized activity 1) emerges from an initial left-biased profile; 2) this left-to-right emergence occurs ∼60-80phd and does not require the presence of a tutor; and 3) also emerges in ZFs raised in a canary auditory environment. Furthermore, lateralization and song development were positively correlated, although these measurements are not necessarily causally related. Awake, bilateral NCM electrophysiology in the same birds when adult, confirmed they were right-lateralized and that lateralized activity for specific test stimuli depended on rearing experience. Lastly, a decoding assay showed that canary-based rearing led to increased decoding accuracy of canary test stimuli, suggesting that neurons exhibit enhanced encodability for those sounds heard earlier. Together, the results document the ontogenetic timeline of auditory lateralization in the songbird and show that auditory experience in development, including passive exposure, shapes how auditory regions in the brain process stimuli in adulthood. Taken together with earlier results showing 1) the absence of lateral differences in ZFs reared in auditory deprivation and 2) that lateralization reverses dynamically in concert with improved discrimination in adult ZFs exposed to novel auditory environments, our current timeline suggests that the dynamic emergence of lateralization reflects the brain’s plastic response to novel auditory experiences.

**Significance Statement:** Lateralized brain processes are ubiquitous in the animal kingdom and are shaped by sensory experience during development. Further, developmental experience can have lasting effects on adult sensory processing. Here, we document, for the first time, the timeline of the emergence of auditory lateralization in the Zebra finch and characterize the lasting effects of rearing experience on adult auditory processing in a higher auditory association region.

## Introduction

Sensory experience early in development – even in the embryo - can have lasting effects on how stimuli are processed in adulthood (Rogers, 1982). A hallmark feature shared by all amniotes’ is the ontogenetic emergence of lateralized behaviors and brain processes (Gunturkun & Ocklenburg, 2017), which requires sensory experience (Rogers, 1982). In general, the ontogenesis of the lateralized brain is driven by genetic factors (e.g. nodal-cascades; Concha et al., 2000) that further interact with sensory and motoric inputs (Chiandetti et al., 2013). At later stages of development, refinement of brain processes can manifest as hemispheric specialization—reflected by lateralized processing of stimuli in adulthood. While lateralization is often studied in the context of human language, for which processing and production is generally proposed to be left-lateralized, neither vocal communication nor hemispheric biases are unique to humans; additionally, lateralized responses likely include activation of areas flanking vocal-processing regions as a function of vocal signal complexity (Peelle, 2012). The main goal of the current study is to reveal the ontogenesis of lateralized brain function in a songbird, the Zebra finch (ZF; *Taeniopygia Guttata*), where both auditory processing and vocal production of communication signals are known to be lateralized in adults.

ZFs are vocal learners (Petkov & Jarvis, 2012) that provide the best-developed animal model for the study of vocal production and processing of communication signals; as in humans (Leybaert & D’Hondt, 2003; Marcotte & Morere, 1990), they require auditory inputs during development in order to display lateralized auditory processing in adulthood (Phan & Vicario, 2010). The songbird higher-order auditory region caudomedial nidopallium (NCM) is a Wernicke-like auditory association region (Moorman et al., 2012) that exhibits the ability to form long-lasting auditory memories in the form of stimulus-specific adaptation (Chew et al., 1995). Similar to areas of the human cortex, NCM exhibits lateralized activity in response to conspecific vocalizations: the right NCM exhibits higher activity in normally-reared adult ZFs, whereas ZFs deprived of auditory input in development are not lateralized (Phan & Vicario, 2010). However, the ontogeny of the right-lateralized NCM profile has not been documented.

Cross-fostering experiments, in which ZFs are raised by a closely-related songbird species, the Bengalese Finch, have suggested that early experiences can shape how vocalizations are encoded in auditory regions (Woolley, 2010; 2012). For example, cross-fostered ZFs exhibit attenuated mutual information (MI) for ZF test stimuli in the songbird primary auditory region (Field L) and midbrain (MLd), although MI decrements could be explained by decreased neuronal firing (Woolley et al., 2010). Behavioral examination of cross-fostered ZFs revealed that their typical preference for conspecific signals is reversed (cross-fostered ZFs prefer songs that are similar to their heterospecific tutor; Clayton, 1990) and that their ability to discriminate conspecific songs diminishes (Campbell & Hauber, 2009); this principle seems to also be shared in wild species of sparrows exposed to songs of a subspecies (Schroeder & Podos, 2023). In addition, ZFs raised by Bengalese finches but maintained in a ZF aviary, display abnormal responsivity, selectivity, and firing consistency to conspecific song, suggesting that social factors can also shape auditory regions regardless of the background auditory environment (Schroeder & Remage-Healey, 2024). Altogether, developmental auditory experience can shape how stimuli are processed and acted upon in adulthood, potentially due to the perceptual boundaries that are shaped by the auditory environment in early life; e.g. in humans, the perception of phonemic boundaries is shaped by early language exposure (Kuhl, 2000). Given that NCM has the ability, via mere exposure (c.f. statistical learning; Saffran et al., 1999), to store statistical information about the environment (Lu & Vicario, 2014; Dong & Vicario, 2020), form auditory memories (Chew et al., 1995), and reflect developmental experience (Phan & Vicario, 2010), it presents a potential locus where different developmental acoustic experiences (e.g. heterospecific exposure) can influence adult auditory processing. Therefore, by revealing the ontogenetic time of auditory lateralization, how it is shaped by early experience and relates to auditory processing in adulthood, we aim to offer domain-specific evidence for mechanisms that reflect developmental principles of brain organization by sampling lateralized processes.

To reveal the ontogenesis of lateralized auditory activity in developing male ZFs and to document the lasting effects of developmental auditory experiences in the adult NCM, young ZFs were removed from their home aviary, socially isolated and assigned to three different auditory rearing conditions: a well-established tutor-driven paradigm, an unfamiliar ZF aviary recording (no tutor),or a heterospecific aviary recording (canary). We report that auditory responses are initially left-biased; typical right-lateralized responses emerge in later stages of development (∼60-80 days post-hatch). In addition, adult NCM neural responses to canary stimuli are better decoded in Can-Env-reared birds than in the Zf-Tut and Zf-Env cohorts. Concomitantly, NCM decoding for ZF test stimuli was lower in Can-Env-reared ZFs. In summary, we provide a time-course of the ontogeny of auditory lateralization in the ZF and offer detailed evidence of lasting effects of developmental experience on adult auditory processing.

## Methods

### Subjects

Young male ZFs were raised with their parents and siblings to 40d post-hatch (phd). They were then removed from their home cage, implanted with an epidural electrode array (described below), and individually housed in acoustical-isolation chambers where they were exposed to one of three auditory conditions (described below): Zf-Tut (n = 7), Zf-Env (n = 8), or Can-Env (n = 7) over the course of development. All birds were maintained on a 12/12-hour daylight cycle (7am-7pm) with food and water accessible *ad libitum*; they remained healthy over the course of the experiment. ZFs were maintained on this regimen until they reached adulthood (defined as120phd), at which point acute NCM electrophysiology was performed. All procedures were approved by the Institutional Animal Care and Use Committee at Rutgers University.

### Rearing Environments

ZFs were raised in three auditory environments to explore whether different rearing conditions would influence the emergence of lateralized biases during development. In addition, this allowed us to test whether these early-life experiences had long-lasting effects in adulthood. Birds were randomly assigned to one of 3 groups. If ZFs originated from the same clutch, they were divided amongst the different exposure conditions. One cohort was reared with an established tutor-driven paradigm (Zf-Tut; Tchernichovski et al., 2000). Briefly, Zf-Tut birds had access to two keys that, when pressed, initiated playback of a tutor song via a speaker (the same tutor song, SAMBA, was used for all Zf-Tut birds). Each day, song playback was available for two sessions (morning and afternoon) and a quota of 20 playbacks was allowed per session (total of 40 possible playbacks per day); key-pressing, tutor-song playback, and song recordings were managed by Sound Analysis Pro (SAP; Tchernichovski et al., 2000).

The other two cohorts experienced environments consisting of daily playback of 12hr recordings from either a novel ZF (Zf-Env) or canary (Can-Env) aviary with no explicit tutoring. These recordings were obtained from aviaries located at the Rockefeller University Field Research Station (Millbrook, NY), which ensured that both were completely novel to the birds at the start of the experiment. The Zf-Env environment consisted of ZF calls and songs, representing a conspecific ‘native’ environment for the subjects; thus, birds exposed to this environment received ethologically-relevant auditory input. The Can-Env environment consisted of calls and songs from canaries, which represent a heterospecific ‘foreign’ environment. If adult auditory processing is shaped by sensory inputs during development, this may be akin to young human infants being adopted into a foreign language environment (which can have an effect on how speech is processed by shaping perceptual boundaries; Kuhl, 2000). For all rearing conditions, omnidirectional condenser microphones were placed in the cages to record and track their song development. Since recording songs from the subjects was impractical in the Zf-Env and Can-Env groups, due to ongoing sounds, auditory playback of the environments was restricted to 10.5 hrs a day (8:30am-7pm); the first 1.5 hrs of everyday (7am-8:30am) consisted of silence, which allowed the birds’ vocalizations to be recorded (via SAP). The incorporation of the 1.5hr silence period was automatically controlled via a subroutine (AutoHotKey v1.1.22) that initiated onset/offset of playback in addition to start/stop of SAP software. All environmental playbacks were set to an average amplitude of 70 dB. **Figure 1** provides visual representations of the acoustic features of all three rearing environments.

**Figure 1.**
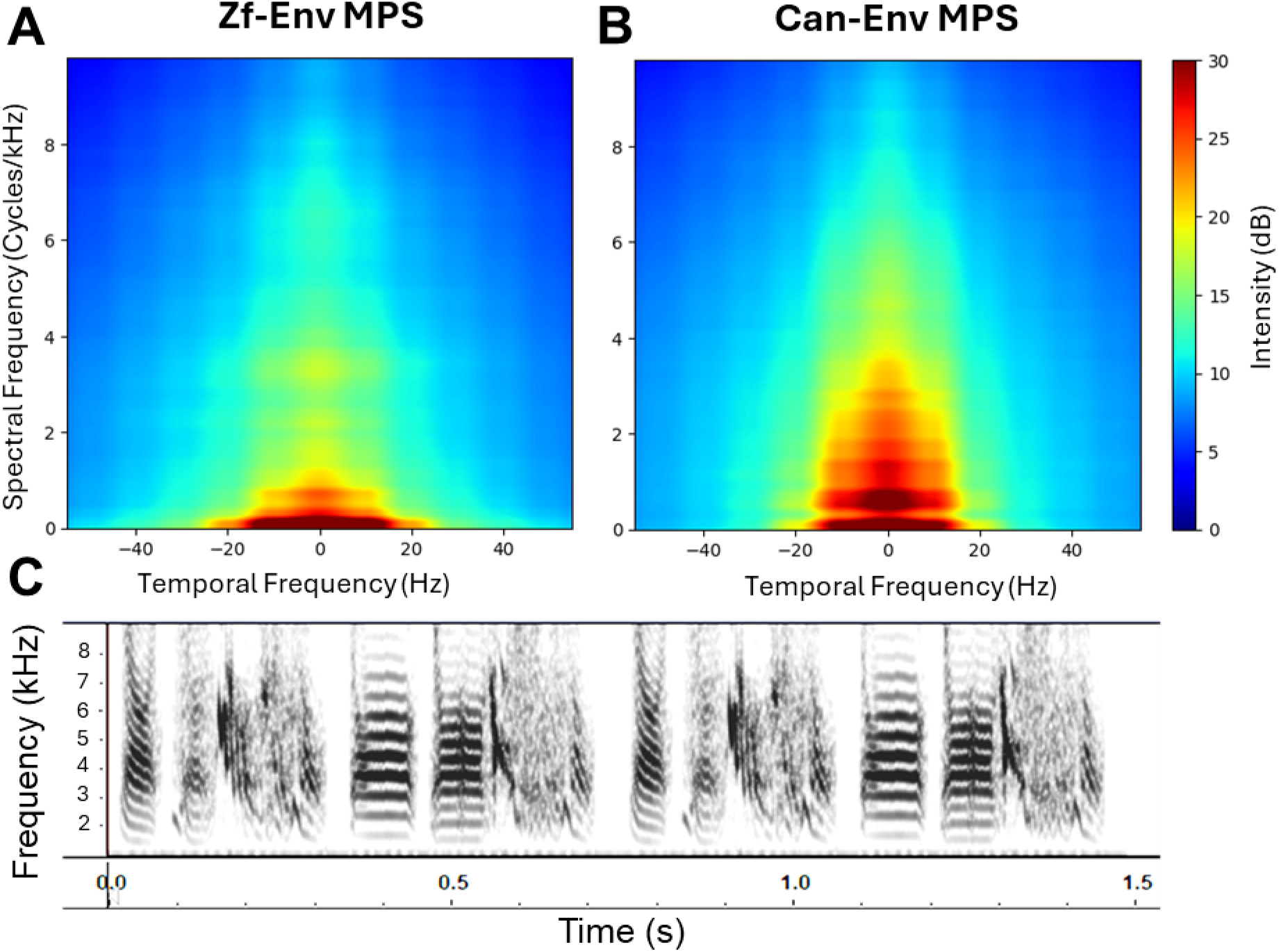
Quantification and visualization of rearing environments. Spectral and temporal modulations of the Zf-Env (A) and Can-Env (B) rearing environments are shown as modulation power spectra (MPS) generated from 15 second samples. For A and B, the y-axes denote the spectral frequency as cycles of power over frequency (Cycles/kHz) and the x-axes denote temporal frequency (Hz); ZFs heard the full 12hr playback of these auditory environments daily throughout development according to their group assignment. C) Sonogram of the tutor song (Samba) with which the Zf-Tut cohort was reared; playback of the tutor song was driven by self-initiated keypecks.

### Surgery

When ZFs reached 40phd, they were removed from their home cage, comfortably swaddled in a plastic tube, anesthetized, and fixed in a stereotaxic apparatus. While under anesthesia (1-3% Isoflurane), the birds’ heads were shaved, a local analgesic was administered (.25% Bupivacaine), and the skull was revealed by creating an incision on the scalp. At 40phd, the skull of the bird is composed of a single layer (adult ZFs’ skulls are composed of two layers connected via spicules), so a small craniotomy was carefully created with a razor blade and the skull around the opening was scored to create a rough surface to which cement would adhere. A head-fixation post was cemented (Anhydrous Polycarboxylate Cement; Tylok Plus, LOT: 161018) on the frontal portion of the skull and a cement well was built around the skull opening.

An epidural array (as described in Furest Cataldo, 2023), consisting of 8 recording pins (2 rows of 4 pins), was lowered into the skull opening and positioned on the dura such that each row of pins flanked the midsagittal sinus (i.e., 4 pins were arranged on the rostro-caudal axis atop each hemisphere). The caudal-most pins were placed near the bifurcation of the midsagittal sinus so that they sat above the auditory forebrain. The epidural array incorporated a connector and a silver ground wire which was tucked between the dura and skull, away from the recording pins; for all birds, the silver wire was tucked towards the left hemisphere, which has been shown to have no effect on lateralized detection of brain activity (Furest Cataldo, 2023). **Figure 2A-B** illustrates the epidural array—a modified integrated circuit chip-holder—and its positioning. Silastic compound (Silicone Elastomer, Kwik-Cast) was used to secure pins’ positions as well as to isolate them from each other. Lastly, cement was used to fix the array to the skull and the previously built cement well. Anesthesia was then discontinued, an analgesic (Meloxicam; 0.01 mL of 5 mg/mL) was administered, and the birds’ recovery was monitored until they could eat and perch. The birds were then returned to their home aviary in an individual cage for 1 day, after which they were individually isolated in their assigned auditory environment.

**Figure 2.**
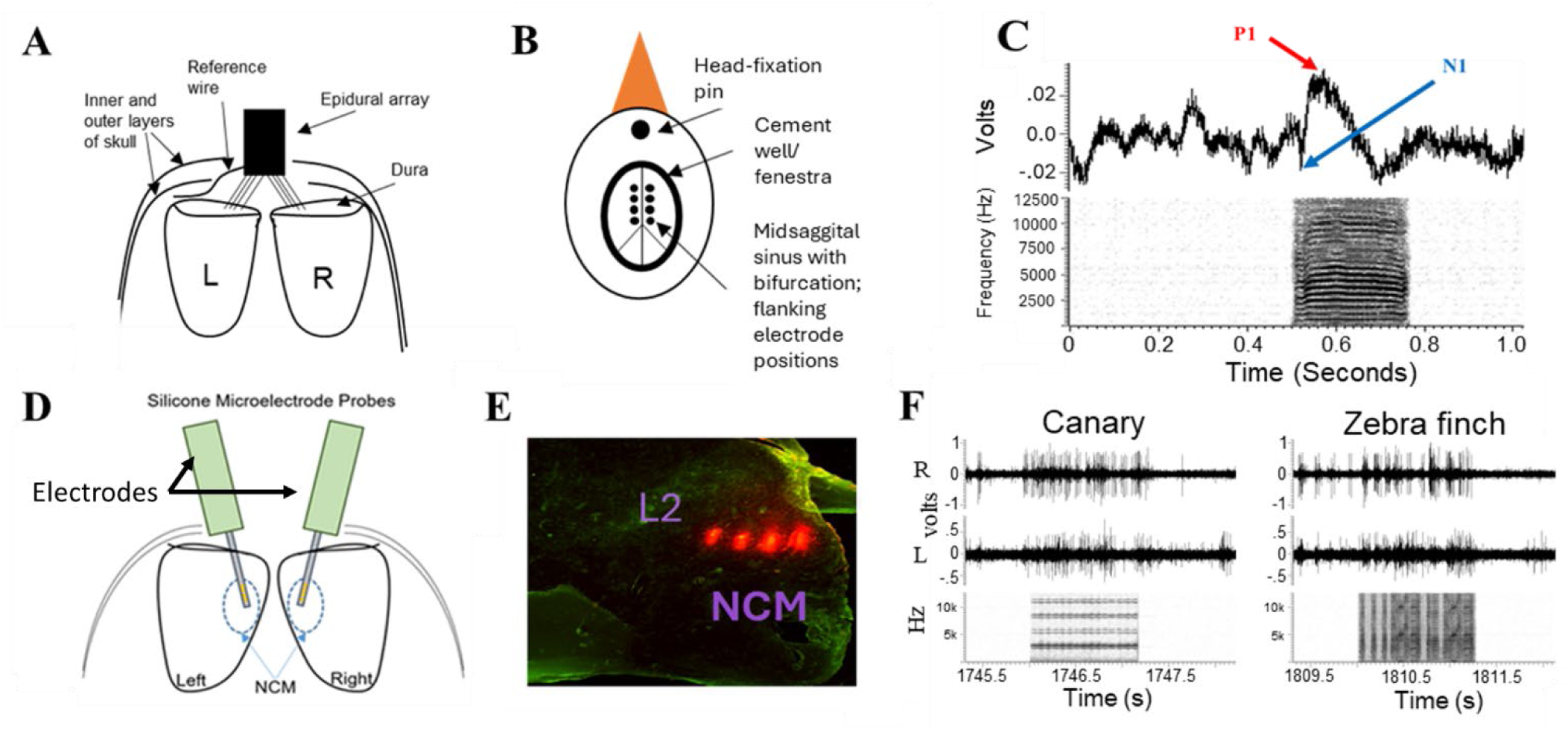
Epidural Electrode array implant, NCM anatomy, chronic and acute electrophysiology. A) An array of 8 electrodes (4 on each hemisphere) were lowered onto the dura. B) The array flanked the midsagittal sinus, with the caudal-most electrodes positioned just rostral to the bifurcation of the sinus. A ground/reference silver wire was tucked between the dura and the inner layer of skull. C) Sample averaged event-related potential (ERP; top) and stimulus sonogram (female call; below) as a function of time (seconds). The magnitude of the ERP was quantified as the peak-to-peak distance (P2P; see methods) between the first positive peak (P1) and first negative peak (N1) of the waveform during the stimulus-on period. D) For acute NCM electrophysiology, silicon multi-electrode probes were inserted bilaterally into NCM. E) Example histology from acute NCM electrophysiology showing electrode traces (red; DiI) in one NCM relative to a nearby structure, Field L2. F) Examples of multiunit activity from each hemisphere in response to zebra finch and canary song stimuli. Panels A-B were adapted from a previous study that employed the same technique (Furest Cataldo et al., 2023).

### Epidural Electrophysiology

A total of 21 epidural recordings were obtained from each ZF across development: recordings took place every 4d from 40-120phd. The first recording was obtained on the day after surgery (when the birds were in their home aviary) and the remaining recordings were obtained after the birds were socially isolated in their respective developmental environments (Zf-Env, Zf-Tut, or Can-Env). On each recording day, the bird was removed from its environment for ∼30mins, comfortably placed in a plastic tube, head-fixed in a stereotaxic apparatus via the head-fixation post, and a miniature 9-channel headstage preamplifier (Neuralynx HS-8-CNR-MDR50) was mated to the implanted epidural array. All recordings were conducted in a large soundproof booth (IAC Inc., Bronx, NY) at a sampling rate of 25kHz. Electrical signals were amplified at a gain of x1000, filtered (bandpass 1-1000Hz), digitized with a Power 1401 A/D converter (Cambridge Electronic Design; CED), and recorded via Spike2 v7.01 software (CED). Bilateral activity from the 8 epidural pins was recorded in response to test stimuli. Each stimulus set consisted of 3 novel female ZF calls (200-300 ms in duration) that were played 100 times each, in shuffled order, with an inter-stimulus interval (ISI; onset-onset) of 5 seconds. Simple female calls were used to reduce the test stimulus’ contribution to the development of lateralized responses; they lack frequency modulations (FM) but elicit quantifiable event-related potentials (e.g. Maul et al., 2010; Furest Cataldo et al., 2023) and have been shown to elicit lateralized NCM activity in normally-reared ZFs (Phan & Vicario, 2010). A different stimulus set was used for each recording time point in each bird to avoid memory effects and the order of stimulus sets was counterbalanced across birds.

### Acute, Multi-unit Electrophysiology

At 118phd, and after the final epidural recording, birds were anesthetized, the entire epidural implant was removed, a new head-fixation pin was affixed to the frontal portion of the skull, and a new cement well was built around the skull opening. Following this surgery, the birds were returned to their rearing booth—with continued access to their acoustic rearing environments—for at least 2 days. Acute, awake bilateral NCM electrophysiology was then performed at 120-124phd by inserting silicon probes (NeuroNexus; 16 sites per probe, 4×4-4mm-200-200-1250-A16 layout; .1-.2 Mohm) into each hemisphere to target NCM (**Figure 2E**); for each hemisphere, insertions were made .5mm lateral of the midsagittal sinus and .5mm rostral from the bifurcation of the midsagittal sinus at a relative depth of ∼1-1.5mm. Before insertion, probes were coated with DiI (10% in ethanol; Sigma Aldrich, St. Louis, MO) to enable histological track reconstruction. Electrical activity was amplified (x10,000), bandpass-filtered (.5-5kHz), digitized with a Power 1401 A/D converter (CED), and recorded via Spike2 v7.01 software (CED); acquisition of activity between the two NCMs was synchronized.

Each probe was lowered while playing a search stimulus (amplitude-modulated band-passed noise, not part of any analysis) until bursting responses characteristic of NCM were observed. Then multi-unit activity was recorded in response to test stimuli (**Figure 2F**). Each stimulus set consisted of 13 stimuli: 5 novel ZF songs, 5 novel canary (CAN) songs, bird’s own song (BOS; specific to each bird), the Zf-Tut tutor song (SAMBA), and the biological father’s song (DAD; specific to each subject). Within a playback set, each stimulus was repeated 25 times in shuffled order with an ISI of 8s. Directly following acute recordings, birds were anesthetized with a lethal dose of Euthasol (pentobarbital sodium and phenytoin sodium; 390mg/mL), transcardially perfused with 0.9% saline followed by 4% paraformaldehyde in PBS and their brains were sectioned to histologically confirm electrode placement.

### Data Analysis: Epidural

Playback of simple stimuli (female ZF calls) elicited changes in electrical activity that were captured by epidural electrodes and characterized as event-related potentials (ERPs). As seen in Figure **2C**, the ERP waveform is composed of a rapid down-going deflection (minimum peak, N1), followed by a rapid upgoing deflection (maximum peak, P1), and ending with a slow ramp-like decrease back to baseline voltage levels. To quantify the ERP magnitude, waveforms were first averaged for each electrode pin separately across the first 25 stimulus presentations of each stimulus. A peak-to-peak (P2P) measure was then calculated by taking the difference between the maximum and minimum peaks of the averaged ERP waveform (i.e. P1-N1; Figure **2C-D**); P2P provided an index of the response magnitude. In order to track, for each bird, the relative activity between each hemisphere as a function of time, a lateralization index (LI; **Figure 2D**) was calculated by computing separate averages of P2P values across the 4 electrodes on the right (R) and left (L) hemispheres respectively, and calculating the quotient between their absolute difference (R-L) and their arithmetic mean (.5*(R+L)).

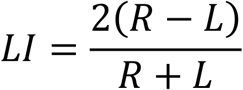

**Equation 1.** The LI is a unitless measure that allows for the comparison between birds and within each bird across development; positive values indicate higher activity on the right (relative to the left) hemisphere and negative values reflect higher activity in the left (relative to the right) hemisphere.

In order to determine the ontogenesis of lateralized activity in the ZF, LIs were tracked for each bird from 40-118phd. LIs in individual ZFs were regressed as a function of time in order to derive the rate of change (slope) and the point at which the regression line crossed LI=0 (zero-int); this allowed the individual differences in the ontogenesis of lateralized auditory activity to be determined. Next, for each rearing condition, LIs were collapsed across individuals to derive the group mean LI change over time. Then, linear regression analysis was performed on these mean LI values to determine the mean LI change over time (slope) and the point at which the mean regression line crossed LI=0 (zero-int) for each condition. In addition, the mean LIs were arbitrarily divided into two half-sessions: the first half of the recordings and the second half of recordings; there were 10 time-point values in the first half and 11 in the second. This allowed for comparisons of the average LI values across two discrete segments of development both within and between groups.

### Data Analysis: Acute Electrophysiology

Prior to any analyses, multi-unit waveforms were thresholded to only include voltage spikes above 2.5 standard deviations from each waveform’s average (as described in Dong & Vicario, 2020). Then stimulus firing rates (FRstim), for each stimulus, trial and electrode, were computed by first summing thresholded spikes during the stimulus presentation (stimulus duration plus 100ms) and dividing it by the sampling duration (stimulus duration + 100ms). Next, baseline FRs (FRb) were computed for each trial by summing the thresholded spikes over the 500ms prior to stimulus onset and dividing it by the sampling duration (500ms). For each electrode, FRbs were then averaged across stimuli in order to derive an electrode-specific 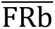, which served to decrease trial-specific noise and enabled an estimate of each site’s average baseline activity. Lastly, for each trial, electrode, and stimulus, the FRb was subtracted from the FRstim (as described in Soyman & Vicario, 2019). This yielded a stable firing rate metric, FR, that is comparable across electrodes and birds since it is controlled for the baseline firing rates for each electrode. FRs were used to compare the magnitude of responding between birds, groups, stimuli, and hemispheres. Lastly, FRs used for comparisons were averaged across the first 2-6 trials of a given stimulus in order to derive a reliable metric of initial response magnitude that is not biased by an electrode site’s adapting response (as in Phan & Vicario, 2010)

In order to determine whether rearing conditions had an effect on stimulus decoding ability in adulthood, thresholded multi-units were first binned into spike-arrays with 10ms bins. Although the stimuli varied slightly in duration, spike-arrays were fixed in size at 1000ms (100 10ms bins). Next, the spike-arrays for each electrode separately were averaged over the first five trials for each stimulus. Lastly, these averaged spike-arrays were z-scored (µ = 0, σ = 1) such that the temporal profile of the responses was normalized for comparison across stimuli. The decoding analysis was based on the 5 ZF and 5 canary (CAN) test stimuli (i.e. BOS, DAD, and SAMBA stimuli were not included). This provided a metric for comparing the ability to decode canary stimuli using multi-unit responses from the different exposure groups. A K-nearest neighbors (KNN) classifier (Pedregosa, 2011) was employed in order to decode the stimulus class (ZF or CAN) that was elicited by the spike trains by using cosine distance a measure of dissimilarity between spike-arrays. This classifier has been previously used to decode sensory activity in crows (Veit & Nieder, 2013) and rodents (e.g. Padmanabhan & Urban 2014). The training dataset was composed of 80% of the responses and the test dataset was composed of the remaining 20%; the composition of the training dataset was stratified such that there was a fair representation of the possible classes. The KNN was employed for each group, bird, and electrode separately; this process was repeated 500 times, with different starting seeds each time (i.e. training/test datasets were different at each seed), and the results were averaged within bird and/or electrode dimensions such that stable classifications could be characterized. The resulting F1 scores, which measure the accuracy of the KNN model based on the precision and recall in the prediction of the spike-train test dataset on stimulus classification, were used as a proxy for decoding accuracy: i.e. the accuracy with which spike-trains could be mathematically decoded into the original stimulus.

### Data Analysis: Song Development

Across development, on each epidural recording day (+/-1 day), three representative song renditions were obtained from each bird. At the conclusion of the experiment, all song renditions were compared to the crystallized version of the respective bird’s song (BOS) obtained at 120phd and to the biological father’s song (DAD) by computing %Similarity scores via SAP (Tchernichovski et al., 2000). For the Zf-Tut birds, %Similarity scores were also computed against the tutor song (SAMBA) they heard in their rearing environment. All %Similarity comparisons were calculated asymmetrically over the song’s time course: “asymmetric” refers to the crystallized song being used as the reference with which to compare each song rendition, and “time course” refers to narrower sections of song being compared between the songs; each song element comparison yielded a %Similarity score which were then combined to produce a global %Similarity score between the two songs. Comparison with BOS assayed the extent to which each finch’s song had stabilized over development at each time-point; i.e. if song development is stable, then %Similarity scores should become higher and vary less during the later stages of development. The comparison against the biological father’s song aimed to detect whether there were residual influences on a given bird’s song over development. Finally, the comparison against SAMBA carried out for the Zf-Tut cohort assayed whether a given bird adopted SAMBA as their tutor song.

For each song sampling day, the three %Similarity values (obtained from the three selected song renditions) were averaged such that each bird had one %Similarity score per sampling day. For each group, %Similarity scores were further collapsed across birds to reveal the average song development per group. Ultimately, these averaged values were employed to determine average bird song development trajectories and to investigate the relationship with changes in lateralized, auditory-evoked, epidural activity.

### Data analysis: General

Statistical analyses were performed using OriginPro (2023b) and included one- and two-way analysis of variance (ANOVA), linear regressions, and post-hoc comparisons using the Holm-Bonferroni method, which controls the family-wise alpha but is more lenient than the Bonferroni correction. All analyses assumed an error rate (alpha) of 0.05. Prediction of the stimulus classes that elicited NCM multiunit activity were carried out by a K-nearest neighbor classifier (Pedregosa, 2011) as described. For visualization purposes, dimensionality reduction of NCM multiunit activity was employed via Uniform Manifold Approximation and Projection (UMAP; McInnes, 2018). Lastly, if equal variances were not assumed in independent-samples t-tests, Welch’s correction was applied. For brevity, factorial ANOVA results for the FR and decoding accuracy analyses are included in **Tables 2.1-2**.

## Results

### Ontogenesis of lateralized auditory-evoked epidural activity in the ZF

To determine the timeline of the emergence of lateralized auditory activity in the developing songbird, young male ZFs were socially isolated in sound-attenuating chambers and exposed to one of three acoustic environments: Zf-Tut, Zf-Env, Can-Env (see Methods). Bilateral epidural activity was chronically tracked in these birds across development. The overall pattern we observed was that most ZFs exhibit left-biased activity during the early part of development, followed by right-lateralized activity later in development and adulthood.

As seen in **Figure 3**, analysis of auditory ERPs recorded with the epidural arrays in developing birds showed the following pattern: LIs tend to be negative (i.e. left-lateralized activity) at early time-points, and progressively change to become positive (i.e. right-lateralized activity). Regression analysis of this pattern of LI change over development showed a significant correlation for all three groups (**Figure 3A-C**): Zf-Tut (Rsq = .78, p <.001), Zf-Env (Rsq = .61, p < .001), and Can-Env (Rsq = .49, p < .001). When all the groups were collapsed, the same pattern was observed (Rsq = .63, p < .01; **Figure 3D**). In addition, the averaged LI values were compared, within groups, between the first (40-80phd) and second (80-120phd) segment of recordings (Supp Figure 2, blue dashed line). For all groups, there were significant differences between the averaged LI values of the first and second segment of recordings: Zf-Tut (t(19) = 6.4, p < .001), Zf-Env (t(19) = 5.25, p < .001), Can-Env (t(19) = 3.22, p = .042). When the group data were collapsed, there was also a significant difference, t(19) = 5.60, p < .001 (Supp Figure 2). Thus these analyses show that ZFs display left-lateralized activity early in development, which gradually changes to the right-lateralized pattern typical of adults.

**Figure 3.**
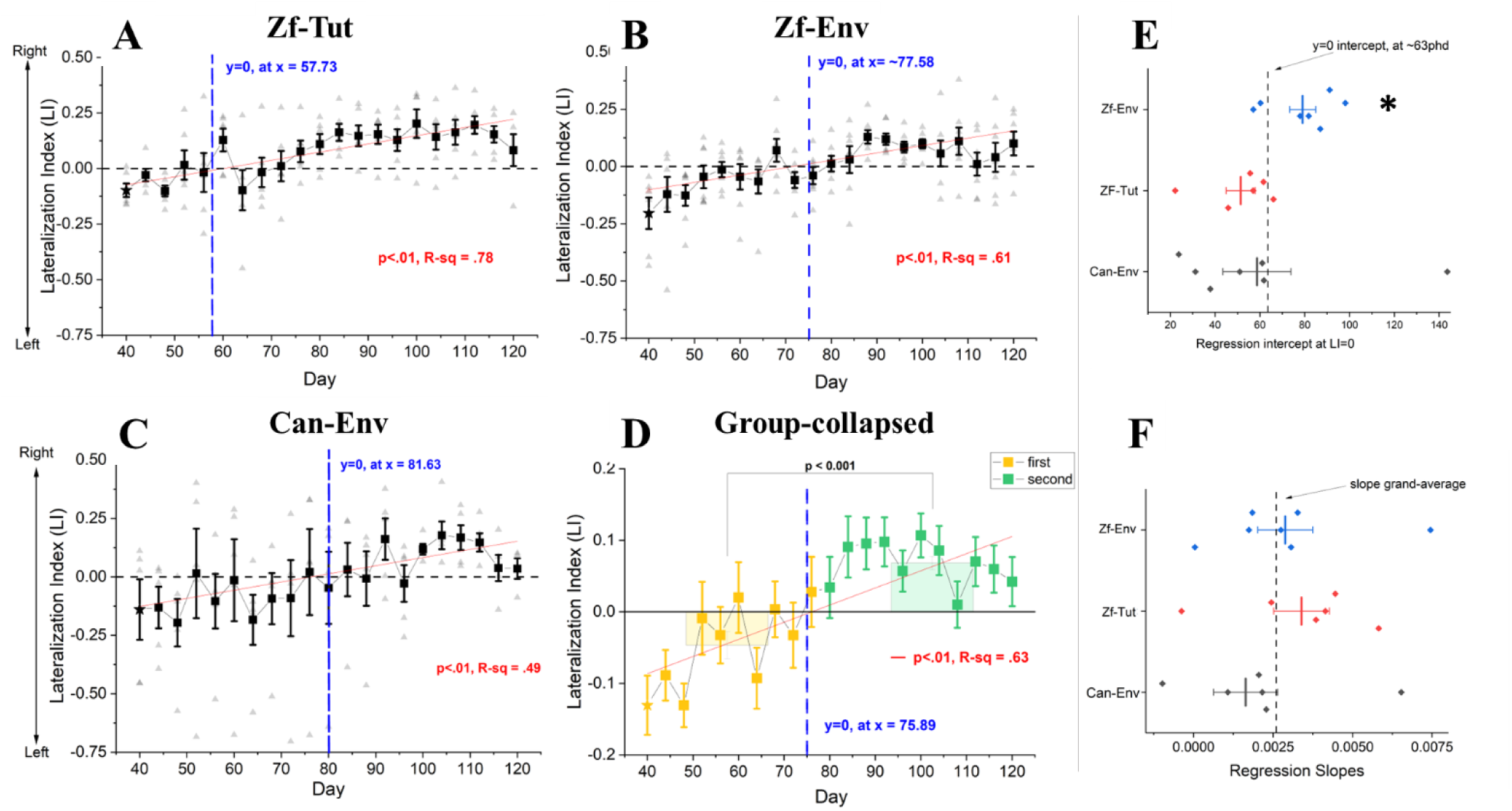
Ontogeny of lateralized epidural activity in development ZFs under 3 different rearing conditions. A, B and C illustrate the mean Lateralization Indices (LI) for each cohort across development (days 40-120); dark squares show means with standard error bars and scattered points show data for individual birds for each testing day. D) Summary of mean LIs across all bird and groups; colored bars show LI values when testing days are divided into first (yellow) and second (green) half-sessions. For A-D, the red lines plot the linear regression for the LI values; the dashed vertical blue lines indicate the y=0 intercept of each regression function. E and F graph the distributions of the y=0 intercept and the LI-regression slopes of all individuals, for each groups, respectively; dashed black lines denote the grand average of the individual metrics. Asterisk in panel E denotes a significant difference in the distribution of y=0 intercepts for the Zf-Env group when compared against the grand-average at an alpha of .05.

The regression analyses were also used to detect whether the onset (zero-int) and rate of LI change (slope) for each individual differed between rearing conditions. Surprisingly, there were no statistical differences between rearing conditions amongst these measures (Figure 3E-F), and only the Zf-Env-reared group exhibited marginally significantly different zero-int scores (t(6) = 2.66, p = .04) relative to the 0-int grand average (derived by averaging all of the zero-int values from every bird in each group; **Figure 3E**). The latter difference would suggest that the time point at which lateralization becomes more right-biased is delayed at the group-level (i.e. occurs later in development, ∼72-83 phd) in the Zf-Env group.

### Typical, adult right-lateralized NCM activity observed for all conditions

As previously established, tutor-reared ZFs display right-biased NCM activity in response to conspecific song playback (Phan & Vicario, 2010). However, the effect of complex acoustic rearing-environments that don’t include explicit tutoring on NCM lateralization have not been previously tested. As a final test of lateralization, we assayed the same ZFs in adulthood via acute, bilateral NCM electrophysiology during song playback. On average, these adult ZFs displayed the typical right-biased pattern of NCM activity for all 3 conditions, which confirmed the right-bias seen in epidural activity sampled during the last epidural recording.

An ANOVA was carried out to determine the extent to which lateralized FRs are affected by rearing conditions (**Figure 4A**). Surprisingly, the different rearing environments did not have a significant effect on the onset, magnitude, or the rate of lateralization change throughout development. Both rearing-group and hemisphere significantly loaded onto the model (group, F(2,2541) = 5.92, p = .003; hemisphere, F(1, 2541) = 63.65, p < .0001); but the group-by-hemisphere interaction was not significant F(2, 2541) = 1.345, p = .26). Post-hoc analysis revealed that, on average, ZFs displayed higher activity in the right hemisphere, relative to the left, in all groups: Zf-Tut (t(2541) = 3.422, p = .0003), Zf-Env (t(2541) = 3.422, p < .0001), Can-Env (t(2541) = 3.422, p < .0001). In addition, there was a positive and significant correlation (**Figure 4B**) between the last epidural LI (sampled at ∼120phd) and an LI derived from NCM multiunit activity (sampled at >120phd), R-sq = .415, p = .005. Overall, adult ZFs displayed right-lateralized NCM activity regardless of rearing condition and the direction of lateralized NCM activity generally agreed with the snapshot of lateralized activity obtained from the last epidural recording.

**Figure 4.**
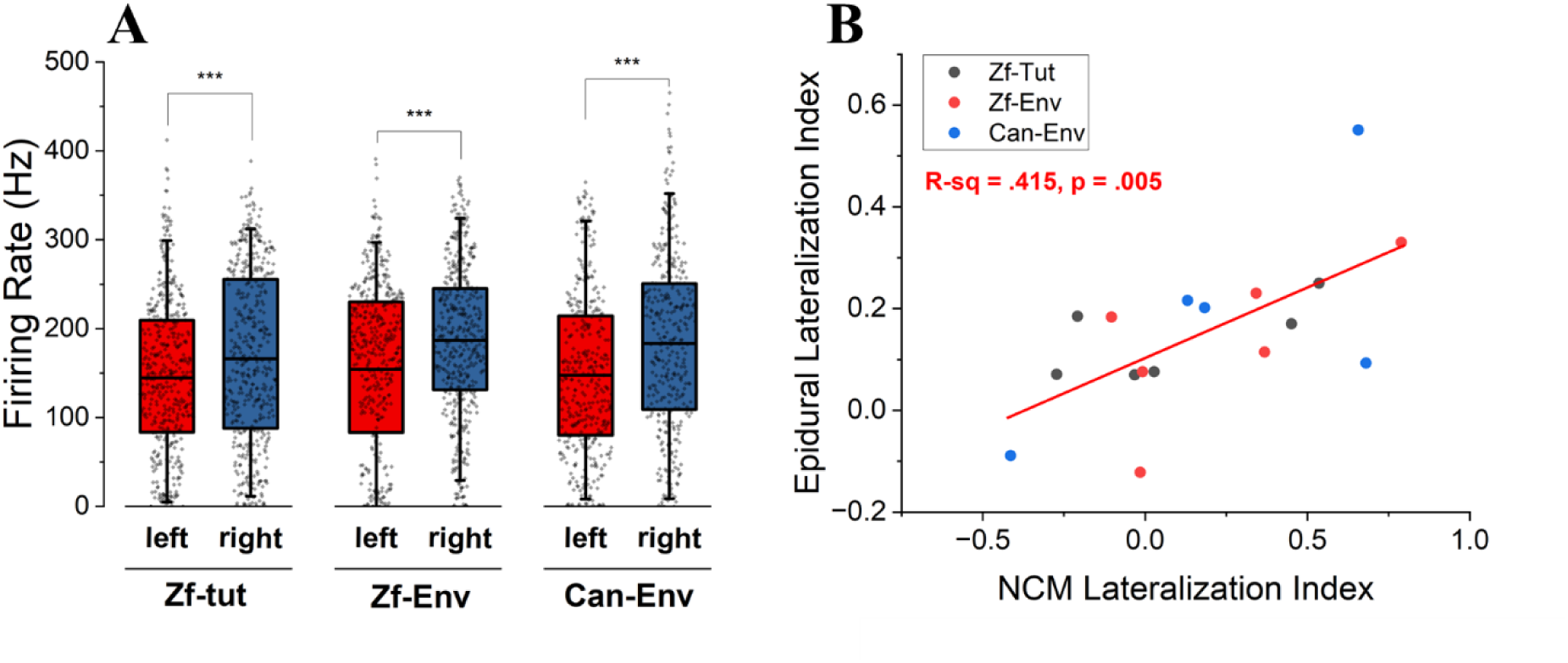
Acute, NCM multiunit firing rates in adulthood as a function of rearing environment and hemisphere. A) All groups displayed higher activity in the right NCM than in the left. B) At the individual level, there was a positive and significant relationship between lateralization indices obtained at the last epidural recording (∼120phd) and the multiunit NCM activity in adulthood (>120-124phd). (***) denotes significance at the .001 alpha level.

### Rearing Environment Effects on Stimulus Processing

In normally-reared ZFs, auditory responses in NCM, measured via electrophysiology or IEG (Zenk) expression, are stronger for conspecific than heterospecific song playback (conspecific bias; Chew et al., 1995; Mello et al., 1992; Yang & Vicario, 2015); e.g., heterospecific canary stimuli elicit lower immediate early gene (IEG) expression in ZFs relative to conspecific Zebra finch stimuli (Mello et al., 1992). Furthermore, earlier observations suggest that naïve, normally-reared ZF adults show relatively more lateralized responses to conspecific than to heterospecific stimuli (Yang & Vicario, 2015). Comparisons of stimulus-elicited FRs between hemispheres, for each rearing group, are illustrated in **Figure 5**. While all groups displayed typical right-lateralized NCM activity in adulthood (**Figure 4A**), further analysis suggested that right-lateralized activity was driven by specific test stimuli with responses that differed depending on rearing group.

**Figure 5.**
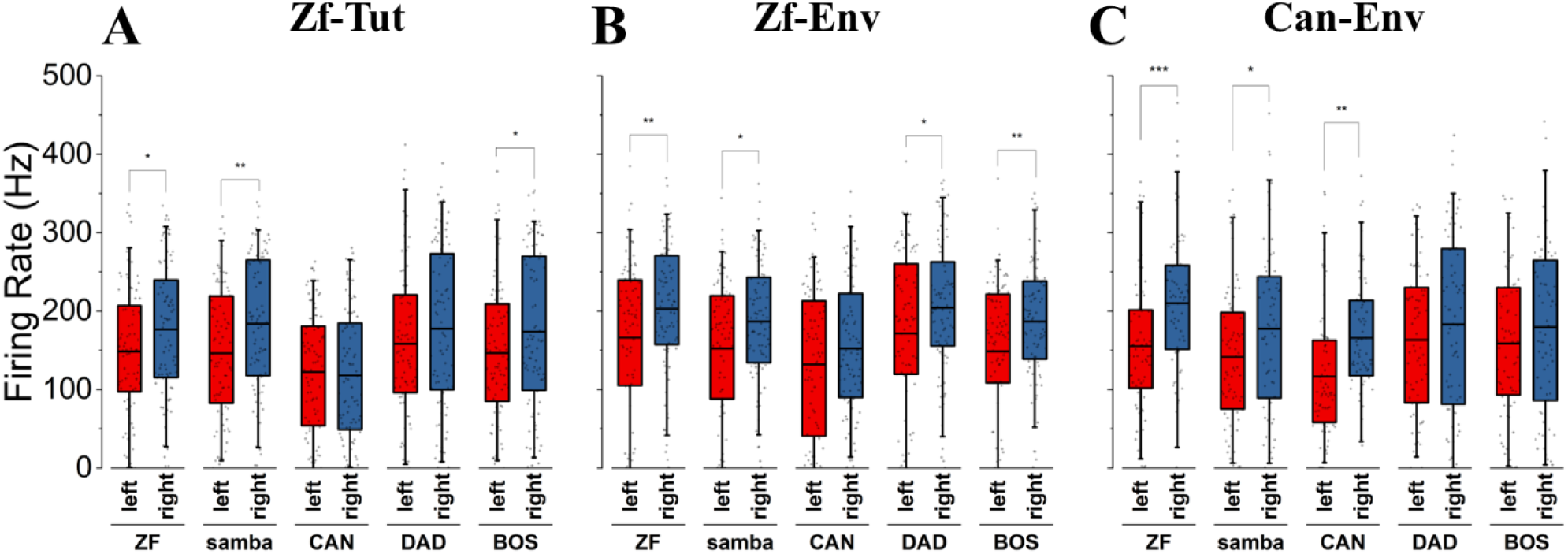
Acute NCM multiunit firing rates as a function of rearing environment, hemisphere, and test stimulus. Firing rates are shown for A) Zf- Tut-reared, B) Zf-Env-reared, and C) Can-Env-reared cohorts. Importantly, zebra finch songs (ZF) elicited right-biased activity in all groups, whereas canary test stimuli (CAN) elicited right-biased activity only in Can-Env raised birds. (*), (**), (***), denote significance at the .05, .01, .001 alpha level, respectively.

A factorial ANOVA with stimulus type (ZF vs CAN) and hemisphere as factors (and the interaction term), conducted for each rearing environment separately, to determine whether specific stimuli were driving the right-biased NCM activity observed in the previous analysis. As seen in Table 1, for all groups, significant main effects were detected for the hemisphere and stimulus factors, but their interaction terms did not reach significance. Post-hoc comparisons were only considered between left and right responses to each stimulus independently. Based on this, the analysis suggested that the hemisphere main effect was driven by different stimuli across all rearing environments. For example, while the ZF and SAMBA stimuli elicited significantly higher activity in the right, relative to the left, NCM, only the Can-Env-reared birds showed similarly right-biased NCM activity in response to canary song stimuli. **Table 1** summarizes the conducted ANOVAs and includes only the significant post-hoc detected via the Holm-Bonferroni method.

**Table 1.**
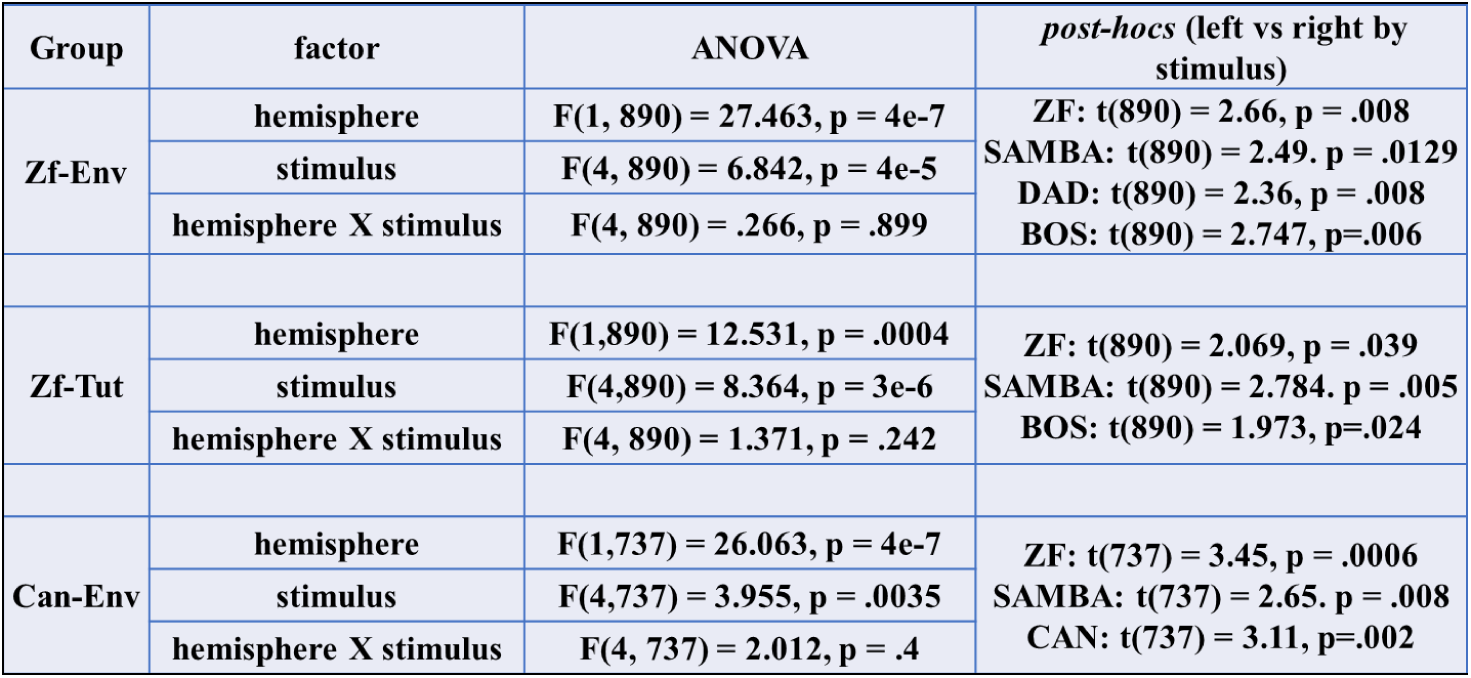
Anova and *post-hoc* comparisons based on NCM electrophysiology performed in ZFs that underwent different rearing conditions (Zf-Env, Zf-Tut, and Can-Env) and playback of different testr stimuli (ZF, CAN, samba, DAD, and BOS songs). ANOVA factors include hemisphere, stimulus, and the interaction term. *Post-hoc* comparisons were conducted, for each group and stimulus separately, across the hemisphere levels.

### Improved decoding capacity for novel canary stimuli in Can-Env birds

Our experiment tested the idea that songbirds reared in different environments, specifically ZFs raised by conspecific (Zf-Tut and Zf-Env) versus heterospecific sounds (Can-Env) will show differences in how stimuli are processed in adulthood. We used a decoding accuracy approach, whereby multi-unit response profiles elicited by either ZF (5 stimuli) or canary (5 stimuli) were mathematically classified and compared to their true category via a K-nearest neighbor classifier (**Figure 6**; see Methods); F1-scores were used as a proxy for decoding accuracy. If passive learning can improve processing of novel sounds by mere exposure (Soyman & Vicario, 2019) and developmental experience can shape how communication signals are processed in adulthood (Kuhl, 2000), one would expect canary-stimulus decoding accuracies to be increased in ZFs in the Can-Env group and not the other groups.

**Figure 6.**
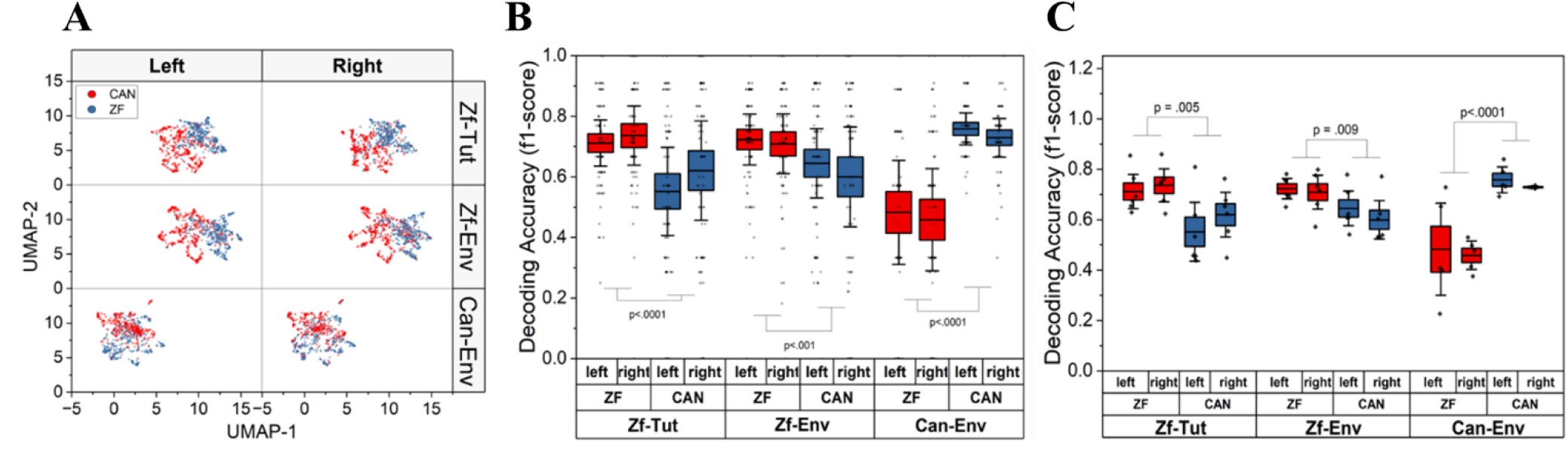
Inspection and classification of stimulus-elicited NCM multiunit activity. A) Visualization of dimensionality-reduced multiunit activity in UMAP-space for canary test stimuli (red circles)y and ZF test stimuli (blue circles). For visualization purposed, stimulus-elicited responses of NCM neurons were plotted in dimension-reduced UMAP-space for each group separately. Each data point represents an electrode channel. B) Decoding accuracy of NCM multiunit activity in the classification of test stimuli, by group and hemisphere. Each data point denotes the accuracy (f1-score) of the employed KNN classifier for each electrode included in the analysis; boxplot boundaries were set to 1 * s.e. of the mean and the whiskers denote 2 * s.e. from the mean. C) Decoding accuracy of NCM multiunit activity in the classification of test stimuli, down sampled for each bird, i.e. each data point represents a bird’ decoding accuracy score by hemisphere and group; boxplot boundaries were set to 1 * s.e. of the mean and the whiskers denote 2 * s.e. from the mean. For all analyses (B and C), the stimulus factor of a two-way ANOVA loaded significantly, suggesting that the variance of the model was best explained by which stimulus was being classified by the KNN; EXP and IMP cohorts displayed higher decoding accuracies for ZF stimuli while the HET cohort displayed higher decoding accuracies for the CAN stimuli. P-

Analysis of decoding accuracy showed that Can-Env-reared ZFs had a heightened ability to distinguish canary test stimuli relative to Zf-Env- or Zf-Tut-raised cohorts. A two-way factorial ANOVA was carried out for each group separately, with stimulus (ZF or canary stimuli) and hemisphere (left or right) as factors; DAD, BOS, and SAMBA stimuli were not used in this analysis. As seen in **Table 2**, for all rearing conditions, neither the hemisphere or interaction terms were significant. However, the stimulus factor loaded significantly into each model, suggesting that, in all groups, decoding accuracies were significantly different between the ZF and canary stimuli in all groups (**Figure 6B**). Interestingly, the direction of the main effect was different depending on rearing condition. That is, the Zf-Tut and Zf-Env reared birds showed higher decodability for ZF songs, relative to canary vocalizations, whereas the opposite pattern was observed in the Can-Env-reared finches. This may be inconsistent with the expectation that there would be a robust innate ability to classify conspecific sounds, although it conforms with behavioral assays conducted in cross-fostered ZFs (Campbell & Hauber, 2009). In addition, while one may have expected that decoding may have been lateralized, given the lateralized FRs observed in Figure 5, this was not observed. However, the results are consistent with the idea that early auditory experience can shape how sounds are processed in adulthood, given that Can-Env-reared finches displayed higher decoding accuracy for canary stimuli.

**Table 2.** ANOVA results, for each rearing condition (Zf- Env, Zf-Tut, and Can-Env), of decoding accuracies as a function of constituent factors: stimulus type (ZF vs CAN), hemisphere (left vs right), and the interaction term (stimulus X hemisphere).

| Group | Factor | ANOVA | **Table 2.** ANOVA results, for each rearing condition (Zf-Env, Zf-Tut, and Can-Env), of decoding accuracies as a function of constituent factors: stimulus type (ZF vs CAN), hemisphere (left vs right), and the interaction term (stimulus X hemisphere). |
| --- | --- | --- | --- |
| Zf-Env | Stimulus | F(1, 352) = 15.449, p = 0.0001 |  |
|  | Hemisphere | F(1, 352) = 1.537, p = 0.21 |  |
|  | Stimulus X Hemisphere | F(1, 352) = 0.412, p = 0.52 |  |
| Zf-Tut | Stimulus | F(1, 354) = 29.936, p = 8e-8 |  |
|  | Hemisphere | F(1, 354) = 3.422, p = .06 |  |
|  | Stimulus X Hemisphere | F(1, 354) = 0.770, p = 0.38 |  |
| Can-Env | Stimulus | F(1, 294) = 114.87, p = 2e-16 |  |
|  | Hemisphere | F(1, 294) = 1.107, p = 0.29 |  |
|  | Stimulus X Hemisphere | F(1, 294) = 0.01, p = 0.92 |  |

To further investigate stimulus-decoding at the individual level, decoding accuracies were collapsed across hemisphere for each individual (**Figure 6C**). Then factorial ANOVAs, with stimulus, hemisphere, and their interaction, were conducted for all rearing groups separately. Similar to the previous analysis, only the stimulus factor loaded significantly: Zf-Tut (F(1, 20) = 10.055, p = .0048), Zf-Env(F(1, 20) = 8.435, p = .009), Can-Env (F(1, 16) = 30.337, p = 4e-5). Again, the direction of the stimulus effect depended on rearing condition, such that Zf-Env- and Zf-Tut-reared ZFs better decoded novel zebra finch (t(20) = −2.904, p = .0088; t(20) = −3.171, p = 0048; respectively), relative to canary, stimuli; again, the opposite was observed for the Can-Env-reared ZFs (t(16) = 5.508, p = 4e-5). In summary, both levels of analyses revealed the same pattern of results: ZFs that were reared in environment that contained conspecific (i.e. zebra finch) acoustics (i.e. Zf-Tut and Zf-Env) showed higher decoding ability for novel zebra finch stimuli and ZFs reared in an environment that contained canary acoustics (i.e. HETNV) had a showed higher decoding ability for novel canary stimuli than for novel zebra finch stimuli (**Figure 6**).

### Song Development and Auditory-evoked Lateralization

Due to the lack of an explicit tutor song in the Zf-Env and Can-Env environments, song development in each bird was assayed via three different comparisons: against crystallized BOS, the biological father’s song (DAD), and SAMBA (for the Zf-Tut group only). For all groups, the %Similarity to DAD across development remained fairly constant throughout development (∼%60-70). This would suggest that, while there may have been residual influences of the father’s song, it was not fully adopted by the birds. Surprisingly, %Similarity to SAMBA remained consistently low (∼%40-50) in the Zf-Tut cohort, despite it being the explicit tutor song (Supp. Figure 3). Lastly, and in order to compare song development across all groups, when each birds’ song renditions across development were compared against their respective crystallized adult BOS, %Similarity dramatically increased as a function of development to resemble BOS more and more as they reached adulthood (**Figure 7A**). While this pattern held true for all groups, Can-Env-reared finches reached only ∼80% %similarity against their final BOS, on average, somewhat lower than scores reached by the Zf-Tut- (∼95%) and Zf-Env-reared (∼93%) finches; the final %Similarity score was derived by comparing BOS obtained at 118phd and the crystallized version at >120phd . In addition, the standard error in %Similarity scores was compared, for each group, as a function of the first and second half of development. The Zf-Tut and Zf-Env finches significantly decreased their standard error in %Similarity (t(14.77) = 3.25, p = .004; t(2.14) = 2.14,p = .046; respectively, equal variances not assumed), while the Can-Env cohort exhibited a relatively stable standard error, indicating similar variability between the two halves of development (t(11.306) = .97, p=.35 **Figure 7B**).

**Figure 7.**
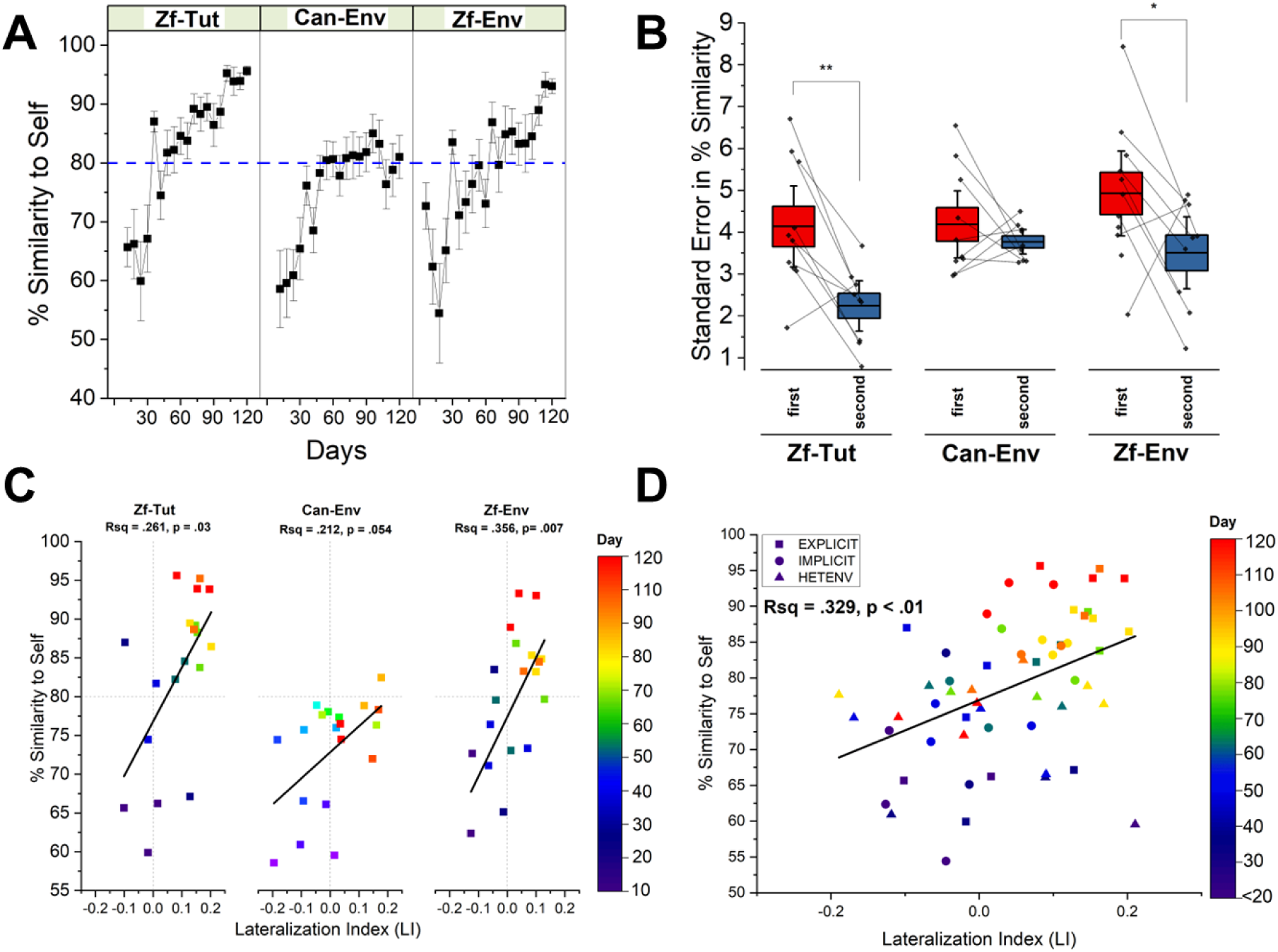
Song development and the ontogenesis of auditory-evoked lateralization. A) Bird-collapsed %Similarity-to-self (as a percent of final adult crystallized song) for each group across time. Zf-Tut and Zf-Env groups reached %Similarity scores that approached 100% (as expected based on the calculus) whereas Can-Env-reared birds stayed around an 80% asymptote (dashed blue line). B) standard errors for %Similarity scores for each group, are subdivided into the first (sessions (T0-T10, i.e. 40-60phd) and second half-sessions (T11-T20, i.e. 80-120phd). Significant differences between sessions were only detected for Zf-Tut and Zf-Env groups. Lines connect data points, between each half-session for each bird. C) Regression analysis between %Similarity scores and Lateralization Index for each group. This analysis was performed independent of time; however, timepoints are indicated by marker colors. D) Similar to (C), but all groups were included in the regression analysis (marker shape denotes rearing group). Partial correlation analysis revealed that the two measurements become uncorrelated when time is included as a controlling variable. (**) and (*) denote significance at the .01 or .05 alpha level, respectively.

Together, ZFs raised in native-like environments (i.e. Zf-Tut and Zf-Env environments) produced songs over development that were more stable (defined by higher %Similarity, and decreased standard errors, to crystallized BOS), whereas ZFs raised in the unfamiliar/foreign Can-Env reached stability in song but low %Similarity-to-self scores; this discrepancy may be explained by the birds being “primed” for a ZF-related environment by their earlier exposure (prior to 40 phd) to ZFs.

In order to reveal the relationship between changes in lateralized epidural activity and song development, group-averaged LIs and %Similarity scores against BOS were regressed independent of time. As seen in Figure 7C, the regressions were significant for the Zf-Tut (Rsq = .261, p = .03) and Zf-Env-reared finches (R-sq = .356, p = .007), and marginally non-significant for the Can-Env-reared birds (R-sq = .212, p = .054). In addition, these relationships do reflect developmental time (color encodes time in **Figure 7C**), both variables seem to correlate over development: lower %Similarity scores (darker colors), which occurred at early stages of development, associated with negative (left-lateralized) LI scores; higher %Similarity scores, at later stages of development, were associated with positive (right-lateralized) LI values. This correlation of %similarity with LI was significant when all groups were collapsed (R-sq = .329, p < .001), and also exhibited the association with developmental time (**Figure 7D**). However, for all groups, the correlation between the two measures dissipates when a partial correlation analysis is conducted with time as a controlling variable: Zf-Tut (r(16) = -.23, p = .34746), Zf- Env (r(17) = -.136, p = .57878), Can-Env (r(16) = -.0005, p=1). This would suggest that the correlation between LI and %Similarity scores is unlikely to be causal, rather both processes likely develop over time but are nevertheless changing with a similar progression. While the onset of lateralization for speech in humans is debated (e.g. Sato, Sagobe, & Mazuka, 2010), it is generally agreed that it arises as a function of speech development and/or proficiency (Holland et al., 2001; Sato, Sogabe, & Mazuka, 2010; Zhang, Gervain, & Roeyers, 2022), a pattern that appears to also be present in the current experiment’s ZFs.

## Discussion

### General

Our results track the ontogenesis of auditory-evoked lateralized epidural activity in developing male ZFs and show the extent to which different rearing environments can shape sensory processing in adulthood. Regardless of rearing environment, all ZFs initially exhibited left-biased epidural activity followed by a shift to right-biased activity later in development that persisted in adulthood. The onset of right-biased activity ranged from ∼60-80phd and differed numerically, but not statistically, between the rearing environments (Figure **2****.3A-C**); the rate at which lateralized responses changed over development was also similar across all rearing conditions. Birds in the Zf-Tut group displayed more robust right-biased patterns of activity in the second half of development, relative to the first half, and compared with the Can-Env and Zf- Env groups (not shown). Chronic epidural recordings enabled us to collect lateralization data longitudinally in individual birds but reflect activity over a region of the caudal forebrain. Our adult electrophysiological data, collected with microelectrodes inserted into NCM of the same birds as adults, confirmed higher activity in the right NCM than in the left in all three groups (**Figure 4A**).

We also showed an effect of rearing environment in adult responses . Canary stimuli elicited significant right-lateralized NCM activity only in in the Can-Env group (**Figure 5**), suggesting that canary sounds had acquired the lateralization pattern typical of conspecific ZF sounds (in normally reared birds) through developmental exposure. In addition, the Can-Env group was distinguished by the ability of NCM neurons to decode canary stimuli in adulthood, as shown by the analysis of decoding accuracy (**Figure 6**). Taken together, the results show that the ontogenesis of lateralization progresses from an initial left-bias to right-bias over the span of development. Furthermore, this change does not depend on hearing conspecific signals (at least after 40phd), and that developmental experience with heterospecific sounds can shape how the adult auditory system processes sounds that are otherwise foreign to the ZF (e.g. Woolley, 2012).

### Ontogenesis of Lateralization in Complex Auditory Environments

The current experiment reared ZFs in three different auditory environments. The Zf-Tut group was reared via a well-established tutor-driven paradigm (Tchernichovski et al., 2000), which has been previously shown to be sufficient to drive right-biased activity when ZFs are tested in adulthood (Phan & Vicario, 2010). However, NCM activity is not lateralized if ZFs are devocalized and reared in silence (Phan & Vicario, 2010), which suggests that auditory exposure to complex communication signals (or at least one, the tutor song) is required to develop typical lateralization patterns. Zf-Env and Can-Env are noisy acoustic spaces with many individuals singing and calling. Thus, these are complex auditory environments, but provide no opportunity to trigger an explicit tutor song, in contrast to Zf-Tut. Despite that, the birds were exposed to multiple examples of either ZF or canary songs in the two environments respectively. Birds in all three environments showed a similar pattern of lateralization changes across development and all were right-lateralized in adulthood. While this is consistent with a role for complex sounds, as previously shown, it further demonstrates that even heterospecific (canary) vocalizations are sufficient for the development of normal lateralization. We may speculate that sufficient exposure to complex speech sounds (of whatever language) may be similarly essential for the development of typical patterns of lateralization in humans.

The results we obtained - an initial left bias that switches to right-bias across development - may be interpreted together with a separate but related experiment in the laboratory. In adult ZFs exposed to a novel acoustic environment for extended periods, typical right lateralization reverses temporarily to left-bias, then returns to right-bias after a couple of weeks; these birds then show improved discrimination of the novel sounds (Furest Cataldo et al., 2023). Effectively, all the young birds in the present study were exposed to a novel acoustic environment, and they showed initial left-biased activity that switched to right bias over weeks of exposure. Together with earlier results showing the absence of lateral differences in ZFs reared in auditory deprivation, our current timeline suggests that the dynamic emergence of right lateralization over ontogeny reflects the brain’s plastic response to novel auditory experiences. Taken together, these experiments provide a framework and conceptual hypotheses for studying the development and dynamics of typical and atypical auditory lateralization and its impact on adult sensory processing in an animal model whose vocal learning processes parallel human speech development.

### Rearing Environment Shapes Auditory Processing in Adulthood

Human speech perception reflects filtering by an individual’s perceptual boundaries, which are shaped by early-life exposure to language (Kuhl, 2000). A mechanism that may explain this observation is the extent to which auditory neurons may be tuned to respond better to certain sounds and sound contrasts (e.g. speech-sounds that were heard in development or in recent experience) while concomitantly leading to a loss of discrimination for speech-sound contrasts from other languages that were never heard. This attunement could be characterized as the formation and implementation of ‘perceptual filters’ (Amin et al., 2013; Bao et al., 2013; Miller & Knudsen, 2001), which subsequently guide and facilitate the categorization of newly-learned sound prototypes, and their exemplars, following a ‘perceptual magnet effect’ (Kuhl, 1991; Iverson & Kuhl, 2000) while lowering the ability to categorize speech sounds of unexposed languages. If neurons are selectively processing sounds in such a manner, then this may manifest as heightened ability to represent, decode, and distinguish sounds from the same acoustic space as that experienced in development. Results from the NCM FR analysis were consistent with this prediction: only Can-Env-reared birds displayed right-lateralized activity in response to canary as well as ZF stimuli. These observations are also consistent with humans that display typical (left-)lateralized activity to a second language only when they learn it in early development (Hull & Vaid, 2007). This suggested that preferential responding and greater hemispheric asymmetry to conspecific than to heterospecific vocalizations, typical of conspecific bias, extended to a heterospecific stimulus if the bird had been reared in the respective heterospecific environment.

Indeed, further analysis of NCM neural responses in ZFs raised in Can-Env demonstrated enhanced decoding accuracy for novel canary stimuli, which was not observed in the ZFs in the other two groups, who had never heard canary songs prior to acute electrophysiology (**Figure 6**). Interestingly, decoding accuracy for ZF stimuli in Can-Env-reared finches was low relative to Zf- Env- or Zf-Tut-reared ZFs. Although this seems counterintuitive given that the birds had at least some exposure (from 0-40phd and perhaps *in ovo*) to other ZFs housed in the general breeding aviary in which they were initially raised, this observation agrees with similar observations from cross-fostered ZFs (Campbell & Hauser, 2009). Nevertheless, our results suggest that the acoustic rearing environments can shape the way in which the adult brain processes stimuli. While these results document neural discrimination of canary stimuli in Can-Env birds, we have not demonstrated improved discrimination for heterospecific sounds at the behavioral level in these birds. However, a related study in the laboratory demonstrated that adult exposed to Can-Env for 14 or 21 days learned a novel canary discrimination far more rapidly than birds without an extended period of Can-Env exposure (Furest Cataldo et al., 2023), so we speculate that the same would be true of the birds reared in Can-Env. In addition, in the experiment by Furest Cataldo et al. (2023), adult ZFs exposed to Can-Env displayed a shift to atypical left-lateralization, which may present a recapitulation of developmental plasticity as is suggested by the current cohort of birds initially showing left-lateralized responses when introduced to their respective auditory environments, which were effectively novel to them.

### Lateralization in Humans and Zebra Finches

Lateralized behavioral and brain processes are ubiquitous in the animal kingdom and are generally proposed to provide an ethological advantage (Vallortigara, Rogers, & Biezza, 1999; Rogers, 2004; Rogers, Vallortigara, & Andrew, 2013). In humans, the language faculty is observed to be lateralized in most individuals, yet its ontogenetic timeline is controversial. For example, different sampling methodologies yield different interpretations on the average timepoint at which the brain becomes lateralized for speech processing (e.g. Hirnstein et al., 2013; Potdevin et al., 2023). Another factor associated to this controversy is that language-related lateralization, regardless of sampling age, is reliant on test stimuli (e.g. Hickok & Poeppel, 2015). For example, single phonemes are not sufficient to drive lateralized responses, whereas words, and their combinations into phrases or sentences, lead to detectable differences in lateralization (Peelle, 2012). Similarly, adult ZFs that showed lateralization after developmental exposure to complex sounds did not show lateralized responses to simple stimuli (tones and noise; Phan & Vicario, 2010). This would suggest that lateralization also arises as a consequence of stimulus complexity, and that this effect may generalize across vocal learners. Further, it is suggested that lateralized responses to speech sounds, as a function of complexity, are likely due to the increased recruitment of flanking brain regions (that may serve higher-order language processing functions) in the left hemisphere relative to the right (Peelle, 2012). Together, these methodological considerations give rise to different estimates of the ontogenesis of speech-related brain processes in humans.

Here, we reveal the ontogenesis of auditory lateralization in the ZF. The ontogenetic timeline was derived via recordings of epidural activity in response to female ZF calls. In adult, normally-reared male ZFs, NCM activity is lateralized for both conspecific calls and song (Phan & Vicario, 2010). ZF calls contain harmonic stacks; in contrast simpler single-band tones and band-limited noise (without harmonic structure) are not sufficient to drive lateralized NCM activity in adulthood (Phan & Vicario, 2010). We exploited the ability of female calls to elicit lateralized responses across development.

In the current experiment, ontogenesis of lateralized responses was determined based on epidural activity sampled above the auditory forebrain and presumably flanking regions. While we detected that, when adult, the ZFs displayed typically right-lateralized activity in NCM, we cannot rule out that flanking auditory regions (e.g. *caudomedial mesopalliuim,* CMM; a higher-auditory region) or supramodal processing units (e.g. *caudolateral nidopallium,* NCL; a multimodal higher-order region), also exhibit lateralized tendencies that emerge as a function of development. Therefore, based on this discussion, one may speculate that: 1) complex ZF vocalizations, such as calls or songs, may drive lateralized activity (in the studied region NCM, as well as structures with shared inputs such as CMM and NCL) in an experience-dependent manner, and 2) the emergence of speech lateralization in humans may depend on the exposure to rich/complex language environments that shape speech-specific processing centers (canonically Broca’s and Wernicke’s areas) in addition to flanking regions that are recruited during processing complex speech signals (Peelle, 2012). Future studies should investigate whether lateralized activity emerges across loci of the afferent auditory pathway as is suggested by the current study.

### Song Development and Lateralization Ontogenesis

In humans, the development of lateralization during native language learning and language proficiency are related in time (e.g. Sato, Sogabe & Mazuka, 2010): lateralization becomes more prominent presumably as there are concomitant improvements in speech production and comprehension. In addition, the emergence of lateralization has been proposed to be a byproduct of learning (“learning biases hypothesis”, Minagawa-Kawai, Cristia, Dupoux, 2011). Here, our indirect measures of proficiency were song similarity (%Similarity-to-self) and stability (i.e. intraday variability in %similarity scores) when song bouts produced by each bird across development were compared to its crystallized song in adulthood. This yielded two expectations regarding proficiency at later stages of development: higher %Similarity-to-self scores and higher stability (i.e. lower variability). This was indeed observed for the Zf-Tut- and Zf-Env-reared ZFs but not for the Can-Env-reared cohort. Moreover, the linear relationship between lateralization (LI) and %Similarity is positive as a function of development (**Figure 7C- D**). However, given the loss of correlation we saw when time was accounted for in the analysis, we do not claim that there is a causal relationship between changes in lateralization and song production, simply that the measures change in tandem as a function of development.

## Conclusion

The songbird motor and auditory systems have long been known to be lateralized in adulthood; however, to our knowledge, the ontogeny of these asymmetries has not been directly explored. Here, we use a longitudinal assay in individual male birds to demonstrate how auditory lateralization changes from left- to right-bias across development; concomitantly, song structure crystallizes as lateralization reaches its typical right-biased pattern. The left to right progression may reflect a pattern in which extended exposure to a novel environment first engages the left hemisphere then the right as the brain encodes the novel acoustic features, a sequence that parallels the changes observed in adults (Furest Cataldo et al., 2023). In addition, we show that novel acoustic experience during development not only drives the ontogeny of lateralization, but can also have a lasting impact on adult auditory processing. Taken together, these experimental results provide a framework and conceptual hypotheses for studying the development and dynamics of typical and atypical auditory lateralization and its impact on adult sensory processing in an animal model whose vocal learning processes parallel human speech development.

## References

Amin, N., et al. (2013). “Selective and efficient neural coding of communication signals depends on early acoustic and social environment.” PLoS One 8(4): e61417.

Bao, S., et al. (2013). “Emergent categorical representation of natural, complex sounds resulting from the early post-natal sound environment.” Neuroscience 248: 30–42.

Campbell, D. L., & Hauber, M. E. (2009). Cross-fostering diminishes song discrimination in zebra finches (Taeniopygia guttata). Animal cognition, 12(3), 481–490. 10.1007/s10071-008-0209-5

Chew, S. J., Mello, C., Nottebohm, F., Jarvis, E., & Vicario, D. S. (1995). Decrements in auditory responses to a repeated conspecific song are long-lasting and require two periods of protein synthesis in the songbird forebrain. Proceedings of the National Academy of Sciences of the United States of America, 92(8), 3406–3410. 10.1073/pnas.92.8.3406

Chiandetti, C., Galliussi, J., Andrew, R. J., & Vallortigara, G. (2013). Early-light embryonic stimulation suggests a second route, via gene activation, to cerebral lateralization in vertebrates. Scientific reports, 3, 2701. 10.1038/srep02701

Clayton N S, (1990).Assortative mating in zebra finch subspecies, Taeniopygia guttata and T. g. castanotis. Philos. Trans. R. Soc. Lond. B Biol. Sci. 330: 351–370. 10.1098/rstb.1990.0205

Concha, M. L., Burdine, R. D., Russell, C., Schier, A. F., & Wilson, S. W. (2000). A nodal signaling pathway regulates the laterality of neuroanatomical asymmetries in the zebrafish forebrain. Neuron, 28(2), 399–409. 10.1016/s0896-6273(00)00120-3

Dong, M., & Vicario, D. S. (2020). Statistical learning of transition patterns in the songbird auditory forebrain. Scientific reports, 10(1), 7848. 10.1038/s41598-020-64671-4

Furest Cataldo, B., Yang, L., Cabezas, B., Ovetsky, J., & Vicario, D. S. (2023). Novel sound exposure drives dynamic changes in auditory lateralization that are associated with perceptual learning in zebra finches. Communications biology, 6(1), 1205. 10.1038/s42003-023-05567-7

Güntürkün, O., & Ocklenburg, S. (2017). Ontogenesis of Lateralization. Neuron, 94(2), 249–263. 10.1016/j.neuron.2017.02.045

Hickok, G., & Poeppel, D. (2015). Neural basis of speech perception. Handbook of clinical neurology, 129, 149–160. 10.1016/B978-0-444-62630-1.00008-1

Hirnstein, M., Westerhausen, R., Korsnes, M. S., & Hugdahl, K. (2013). Sex differences in language asymmetry are age-dependent and small: a large-scale, consonant-vowel dichotic listening study with behavioral and fMRI data. Cortex; a journal devoted to the study of the nervous system and behavior, 49(7), 1910–1921. 10.1016/j.cortex.2012.08.002

Holland, S. K., Plante, E., Weber Byars, A., Strawsburg, R. H., Schmithorst, V. J., & Ball, W. S., Jr (2001). Normal fMRI brain activation patterns in children performing a verb generation task. NeuroImage, 14(4), 837–843. 10.1006/nimg.2001.0875

Hull, R., & Vaid, J. (2007). Bilingual language lateralization: a meta-analytic tale of two hemispheres. Neuropsychologia, 45(9), 1987–2008. 10.1016/j.neuropsychologia.2007.03.002

Iverson, P., & Kuhl, P. K. (2000). Perceptual magnet and phoneme boundary effects in speech perception: Do they arise from a common mechanism? Perception & Psychophysics, 62(4), 874–886. 10.3758/BF03206929

Kuhl P. K. (2000). A new view of language acquisition. Proceedings of the National Academy of Sciences of the United States of America, 97(22), 11850–11857. 10.1073/pnas.97.22.11850

Kuhl, P. K. (1991). Human adults and human infants show a “perceptual magnet effect” for the prototypes of speech categories, monkeys do not. Perception & Psychophysics, 50(2), 93–107. 10.3758/BF03212211

Leybaert, J., & D’Hondt, M. (2003). Neurolinguistic development in deaf children: The effect of early language experience. International Journal of Audiology, 42(Suppl1), S34– S40. 10.3109/14992020309074622

Lu, K., & Vicario, D. S. (2014). Statistical learning of recurring sound patterns encodes auditory objects in songbird forebrain. Proceedings of the National Academy of Sciences of the United States of America, 111(40), 14553–14558. 10.1073/pnas.1412109111

Marcotte, A. C., & Morere, D. A. (1990). Speech lateralization in deaf populations: evidence for a developmental critical period. Brain and language, 39(1), 134–152. 10.1016/0093-934x(90)90008-5

Mello, C. V., Vicario, D. S., & Clayton, D. F. (1992). Song presentation induces gene expression in the songbird forebrain. Proceedings of the National Academy of Sciences of the United States of America, 89(15), 6818–6822. 10.1073/pnas.89.15.6818

Miller, G. L. and E. I. Knudsen (2001). “Early auditory experience induces frequency-specific, adaptive plasticity in the forebrain gaze fields of the barn owl.” J Neurophysiol 85(5): 2184–2194.

Minagawa-Kawai, Y., Cristià, A., & Dupoux, E. (2011). Cerebral lateralization and early speech acquisition: a developmental scenario. Developmental cognitive neuroscience, 1(3), 217–232. 10.1016/j.dcn.2011.03.005

Moorman, S., Gobes, S. M., Kuijpers, M., Kerkhofs, A., Zandbergen, M. A., & Bolhuis, J. J. (2012). Human-like brain hemispheric dominance in birdsong learning. Proceedings of the National Academy of Sciences of the United States of America, 109(31), 12782–12787. 10.1073/pnas.1207207109

Padmanabhan, K., & Urban, N. N. (2014). Disrupting information coding via block of 4-AP-sensitive potassium channels. Journal of neurophysiology, 112(5), 1054–1066. 10.1152/jn.00823.2013

Peelle J. E. (2012). The hemispheric lateralization of speech processing depends on what “speech” is: a hierarchical perspective. Frontiers in human neuroscience, 6, 309. 10.3389/fnhum.2012.00309

Petkov, C. I., & Jarvis, E. D. (2012). Birds, primates, and spoken language origins: behavioral phenotypes and neurobiological substrates. Frontiers in evolutionary neuroscience, 4, 12. 10.3389/fnevo.2012.00012

Phan, M. L., & Vicario, D. S. (2010). Hemispheric differences in processing of vocalizations depend on early experience. PNAS Proceedings of the National Academy of Sciences of the United States of America, 107(5), 2301–2306. 10.1073/pnas.0900091107

Potdevin, D., Adibpour, P., Garric, C., Somogyi, E., Dehaene-Lambertz, G., Rämä, P., Dubois, J., & Fagard, J. (2023). Brain Lateralization for Language, Vocabulary Development and Handedness at 18 Months. Symmetry, 15(5), 989. 10.3390/sym15050989

Rogers, L. J., Vallortigara, G., & Andrew, R. J. (2013). Divided brains: The biology and behaviour of brain asymmetries. Cambridge University Press. 10.1017/CBO9780511793899

Rogers, L. J., Zucca, P., & Vallortigara, G. (2004). Advantages of having a lateralized brain. *Proceedings*. Biological sciences, 271 *Suppl 6*(Suppl 6), S420–S422. 10.1098/rsbl.2004.0200

Rogers, L. Light experience and asymmetry of brain function in chickens. Nature 297, 223– 225 (1982). 10.1038/297223a0

Sato, Y., Sogabe, Y., & Mazuka, R. (2010). Development of hemispheric specialization for lexical pitch-accent in Japanese infants. Journal of cognitive neuroscience, 22(11), 2503–2513. 10.1162/jocn.2009.21377

Schroeder K.M., Podos, J. (2023). Early exposure to songs of another subspecies enhances song discrimination in wild sparrow nestlings. Animal Behaviour. (203) 123-132. 10.1016/j.anbehav.2023.06.015

Schroeder, K. M., & Remage-Healey, L. (2024). Social and auditory experience shapes forebrain responsiveness in zebra finches before the sensitive period of vocal learning. The Journal of experimental biology, 227(21), jeb247956. 10.1242/jeb.247956

Soyman, E., & Vicario, D. S. (2019). Rapid and long-lasting improvements in neural discrimination of acoustic signals with passive familiarization. PloS one, 14(8), e0221819. 10.1371/journal.pone.0221819

Tchernichovski, O., Nottebohm, F., Ho, C. E., Pesaran, B., & Mitra, P. P. (2000). A procedure for an automated measurement of song similarity. Animal Behaviour, 59(6), 1167– 1176. 10.1006/anbe.1999.1416

Vallortigara, G., Rogers, L. J., & Bisazza, A. (1999). Possible evolutionary origins of cognitive brain lateralization. Brain research. Brain research reviews, 30(2), 164–175. 10.1016/s0165-0173(99)00012-0

Veit, L., Nieder, A. (2013). Abstract rule neurons in the endbrain support intelligent behaviour in corvid songbirds. Nat Commun 4, 2878. 10.1038/ncomms3878

Woolley S. M. (2012). Early experience shapes vocal neural coding and perception in songbirds. Developmental psychobiology, 54(6), 612–631. 10.1002/dev.21014

Woolley, S. M., Hauber, M. E., & Theunissen, F. E. (2010). Developmental experience alters information coding in auditory midbrain and forebrain neurons. Developmental neurobiology, 70(4), 235–252. 10.1002/dneu.20783

Yang, L. M., & Vicario, D. S. (2015). Exposure to a novel stimulus environment alters patterns of lateralization in avian auditory cortex. Neuroscience, 285, 107–118. 10.1016/j.neuroscience.2014.10.022

Zhang, F., Gervain, J., Roeyers, H. (2022). Developmental changes in the brain response to speech during the first year of life: A near-infrared spectroscopy study of dutch-learning infants. Infant behavior and Development, (67), 101724. 10.1016/j.infbeh.2022.101724

